# *Pex11*β knockdown decreases peroxisome abundance and reverses the inhibitory effect of palmitate on pancreatic beta-cell function

**DOI:** 10.1101/2020.04.03.023416

**Authors:** Helen R Blair, Cara Tomas, Audrey E Brown, Satomi Miwa, Alan Health, Alison Russell, Michael-van Ginkel, David Gunn, Mark Walker

## Abstract

**Aims:** Reactive oxygen species generated by the peroxisomes and mitochondria contribute to lipotoxicity in pancreatic beta-cells. Through targeted *Pex11*β knockdown and peroxisome depletion, our aim was to investigate the specific contribution of peroxisomes to palmitate mediated pancreatic beta-cell dysfunction.

**Methods:** MIN6 cells were transfected with probes targeted against *Pex11*β, a regulator of peroxisome abundance, or with scrambled control probes. Peroxisome abundance was measured by PMP-70 protein expression. 48hrs post transfection, cells were incubated with or without 250μM palmitate for a further 48hrs before measurement of reactive oxygen species, mitochondrial respiratory function, and glucose stimulated insulin secretion.

**Results:** *Pex11*β knockdown decreased target gene expression by more than 80% compared with the scrambled control (P<0.001), leading to decreased PMP-70 expression (p<0.01). *Pex11*β knockdown decreased palmitate mediated generation of reactive oxygen species (P<0.001), but with no effect on mitochondrial respiratory function. At 25mM glucose, palmitate treatment decreased insulin secretion in the control cells (2.54±0.25 vs 7.07±0.83 [mean±SEM] ng/hr/μg protein; P<0.001), with a similar pattern in the *Pex11*β knockdown cells. However, in the presence of palmitate, insulin secretion was significantly higher in the *Pex11*β knockdown versus control cells (4.04±0.46 vs 2.54±0.25 ng/hr/μg protein; p<0.05).

**Conclusion:** *Pex11*β knockdown decreased peroxisome abundance, decreased palmitate mediated ROS generation, and reversed the inhibitory effect of palmitate on insulin secretion. These findings highlight a specific and independent role for peroxisomes in pancreatic beta-cell lipotoxicity.

## Introduction

Increased adiposity is a risk factor for type 2 diabetes. Ectopic fat deposition arises when excess lipid is stored in tissues other than adipose tissue, such as the liver, muscle and pancreas. Lipid excess in the pancreas is associated with pancreatic beta-cell dysfunction and impaired insulin secretion (1–4). In vitro, chronic treatment with the long chain saturated fatty acid palmitate has been shown to reduce insulin secretion in human, rat and mouse islets (5–7) and pancreatic beta-cell lines (7–10).

Fatty acid β-oxidation in animals and man is conducted by both peroxisomes and mitochondria (11, 12). Peroxisomes have a role in β-oxidation of long and medium chain fatty acids, with the intermediate products transported to the mitochondria for complete oxidation (12, 13). The first step of peroxisomal β-oxidation generates reactive oxygen species (ROS), principally hydrogen peroxide (H_2_O_2_). In the majority of cells, this is catabolised by oxidoreductase catalase. However, pancreatic β-cells are essentially deficient in this enzyme (14–16) leaving them at risk of lipotoxicity from excess H_2_O_2_ production. Elsner and colleagues demonstrated that peroxisomes are a major source of H_2_O_2_ following palmitate treatment of insulin secretory cells. Overexpression of catalase in peroxisomes and in the cytosol decreased ROS generation by both the peroxisomes and the mitochondria, and led to an improvement in cell viability (17). It remains to be shown, however, whether targeted decrease in peroxisome ROS generation specifically improves pancreatic beta-cell function in the presence of palmitate.

To address this question, we developed a model of targeted peroxisome depletion in MIN6 cells. Peroxisome proliferation occurs by de novo generation from the endoplasmic reticulum and through the division of pre-existing peroxisomes that involves Pex11 proteins (18–21). Pex11β is a protein responsible for the constitutive turnover of peroxisomes and *Pex11*β knockout mice have decreased peroxisome abundance (22–24). Through siRNA silencing of the *Pex11*β gene we decreased peroxisome abundance and ROS generation in MIN6 cells, and found improved beta-cell function in the presence of palmitate.

## Materials and Methods

### Cell Culture and Palmitate Medium

MIN6 cells were donated by Dr Catherine Arden (Diabetes Research Group, Newcastle University, UK) and experiments were performed with cells of passage 23-30. MIN6 cells were cultured in DMEM containing 4500mg/L glucose, L-glutamine and sodium bicarbonate (Sigma), and supplemented with 15% FBS (Life Technologies), 1% Penicillin/Streptomycin (Life Technologies), and 0.0005% β-mercaptoethanol (Sigma). Cells were incubated at 37°C, 5% CO_2_ and passaged when 70-80% confluent. For relevant experiments, MIN6 cells were incubated with 250μM palmitate conjugated to BSA for the final 48hrs before end point measurements were taken. Palmitate was dissolved in deionised water at 70°C before BSA (dissolved in PBS) was added to give a final 4mM stock palmitate solution with a 5:1 ratio, BSA:Palmitate. The palmitate stock was diluted in supplemented DMEM to the final concentration before treatment.

### Transfection with siRNA against *Pex11*β

MIN6 cells were transfected using the Neon Transfection System (Life Technologies) as previously described (25). Two predesigned Ambion siRNA probes against *Pex11*β, s71497 and s71499 (Life Technologies), and a Scrambled siRNA negative control (Life Technologies) were used for the transfection at a concentration of 100nM. The siRNA sequences for the *Pex11*β probes were as follows: s71497, 5’-UCAUGAAUCUGAGCCGUGAtt-3’; 3’- UCACGGCUCAGAUUCAUGAtg-5’, and s71499, 5’- CAACCGAGCCUUGUACUUUtt-3’; 3’- AAAGUACAAGGCUCGGUUGag-5’.

### Real-time PCR

72hr or 96hr after transfection, RNA was extracted using GenElute™ Mammalian Total RNA Miniprep Kit (Sigma) according to the manufacturer’s instructions. Following quantification of the RNA on a NanoDrop 2000 Spectrophotometer (Thermoscientific), cDNA was synthesised using the High Capacity cDNA Reverse Transcription Kit (Applied Biosystems), according to the manufacturer’s instructions. Real-time PCR was carried out on LightCycler^®^480 (Roche) using SYBR-green. Predesigned QuantiTect^®^ Primer Assays (Qiagen) were used for all genes: *Pex11*β and Ykt6. Ykt6 was used as the reference gene. The PCR reactions were carried out using LightCycler 480 SYBR green I mastermix (Roche). Results were analysed using the comparative C_T_ (ΔΔC_T_) method.

### Immunofluorescence

MIN6 cells grown directly onto sterile cover slips were washed with PBS then fixed with 4% paraformaldehyde for 20 minutes at room temperature. Cells were permeabilised with 0.2% t-octylphenoxy-polyethoxyethanol (Triton x-100, Sigma) for 45 minutes. Coverslips were incubated with 20% FBS in PBS for 1 hour at room temperature to block non-specific binding. Cells were incubated with 1:600 dilution of rabbit anti-PMP-70 (Abcam) in 0.05% FBS in PBS at 20-22°C for 1 hour. Cells were then washed with PBS, and incubated with 1:300 dilution of anti-rabbit Alexa Fluor^®^ 546 (Life Technologies) in 0.05% FBS in PBS for one hour at room temperature in the dark. The cover slips were then washed again in PBS before mounting on slides with Vectashield Mounting Medium with DAPI (Vector). Slides were visualised on a Zeiss LSM 780 confocal microscope. Quantification of the images was calculated as the number of peroxisomes per area of the image that was covered by cells.

### Western Blot

Cells were harvested in protein extraction buffer (100mM Tris-Cl, pH 7.4, 100mM KCl, 1mM EDTA, 25mM KF, 0.1% Triton X-100, 0.5mM sodium orthovanadate, 1x protease inhibitor cocktail (Thermo Scientific)) and sonicated for approximately 10 seconds at 5μm amplitude. Protein concentrations were determined using Coomassie blue, and read on a spectrophotometer at 595nm. lOμg protein was boiled for 5 min with sample buffer (62.5mM Tris-HCl pH 6.8, 2% SDS, 10% Glycerol, 0.002% Bromo-phenol blue, 5% ⍰-mercaptoethanol) before being separated on a 10% SDS-PAGE gel. The separated proteins were electrotransferred onto nitrocellulose membranes before being blocked for 1hr in TBS-Tween (65mM Tris-HCl, pH 7.4, 150mM NaCl, 0.1% Tween) containing 5% Marvel milk powder. Following incubation with the appropriate primary and secondary antibodies in 1% Marvel, detection was carried out with the addition of enhanced chemiluminescent solution (Thermo Fisher Scientific), and exposure of the nitrocellulose membrane to X-ray film. PMP-70 antibody was used at a 1.1000 dilution, and β-actin at a 1:10,000 dilution. Quantification of the protein bands was carried out using a GS-800 Calibrated Densitometer (BioRad) and the BioRad software Quantity One 4.2.3.

### Glucose Stimulated Insulin Secretion (GSIS)

The method for carrying out GSIS was adapted from Ishihara et al. 1994 (26). Cells were starved for 30 mins with Krebs-Hepes buffer (119mM NaCl, 4.74mM KCl, 2.54mM CaCl_2_, 1.19mM MgCl_2_, 1.19mM KH_2_PO_4_, 25mM NaHCO_3_, 10mM Hepes, 0.5% BSA pH 7.4), before being incubated with Krebs-Hepes buffer for a further 1 hour with either 3mM (basal), or 25mM (stimulating) glucose concentrations. Following incubation, supernatants were collected and insulin secretion determined using an Insulin ELISA kit (Mercodia) according to the manufacturer’s instructions. Insulin secretion was normalised to total protein content.

### Insulin Content

Following GSIS with 3mM glucose, cells were washed with PBS and harvested in 50μl dH_2_O before being sonicated. 150μl acid ethanol (0.18 M HCl in 100% ethanol) was added to the samples and sonicated further. Samples were then stored at 4°C for 12 hours before insulin content was analysed by insulin ELISA using at least 1.100 dilution. Insulin content was normalised to total protein content.

### Fatty acid oxidation

96hrs post transfection which included 48hrs palmitate incubation, fatty acid oxidation was assessed by using the Seahorse XF24 Analyzer (Agilent Technologies) to measure mitochondrial respiration. Media was replaced with basic media containing 3% FBS and 2mM L-glutamine with or without 250μM palmitate, and cells were placed in a CO_2_ free incubator for 1hr. Oxygen consumption rates (OCR) were used to assess mitochondrial respiration by measurement before and after the injection of compounds that inhibit different mitochondrial complexes: 1μg/ml Oligomycin to inhibit Complex V (ATP synthase) which inhibits the generation of mitochondrial ATP, 2μM and 3.5μM of carbonyl cyanide p-trifluoromthoxy-phenylhydrazone (FCCP) which uncouples respiration, and finally Antimycin A, a mitochondrial complex III (Ubiquinol-Cytochrome c Reductase) inhibitor; which, by preventing electron transfer, results in the abolishment of ATP synthesis and respiration. Basal mitochondrial respiration = basal oxidation - non-mitochondrial oxidation. ATP synthesis by oxidative phosphorylation = ((Basal oxidation - non-mitochondrial oxidation) - (oligomycin inhibited oxidation - non-mitochondrial oxidation)) multiplied by 2.5 (the established phosphate/oxygen ratio for oxidation of palmitate (27)), and further multiplied by 2 (accounting for the 2 oxygen atoms per oxygen molecule).

### Reactive oxygen species (ROS) detection

96hrs post transfection which included 48hrs palmitate incubation, ROS were detected using 2’,7’ –dichlorofluorescin diacetate (DCFDA). ROS such as hydrogen peroxide, peroxyl radicals, and peroxynitrile anions can be detected in live cells when DCFDA is oxidised by the ROS to form the fluorescent dye DCF (28). Cells were washed with PBS and 20μM DCFDA added. Cells were incubated at 37°C, 5% CO_2_ for 45 minutes. Cells were washed twice with PBS and then incubated in supplemented PBS (90% PBS; 10% FBS) for 30 minutes. Fluorescence was recorded (Ex/Em = 485/535). Results were corrected for protein concentration.

### Statistical Analysis

All statistical analysis was carried out using Graphpad Prism 7 (GraphPad Software, San Diego, California, USA). Data are presented as mean ± SEM (standard error of the mean). Statistical significance was tested through use of One way ANOVA followed by an unpaired t-test. Results were considered to be significant when the probability (p) value was <0.05.

## Results

### Palmitate treatment and cell viability

The first step was to investigate the effects of 250μM palmitate on cell viability and apoptosis in MIN6 cells. Following treatment with 250μM palmitate for 48hrs, LDH release and caspase 3/7 activity were measured as indices of cell viability and apoptosis, respectively. There were no differences in LDH release or caspase 3/7 activity between the palmitate treated and BSA control cells (S Fig 1). Under the same conditions, we showed that insulin secretion at 25mM glucose was decreased by >50% in palmitate treated cells compared with the BSA controls; 2.45±0.41 and 5.27±0.83 ng insulin/μg protein respectively (P<0.05) (S Fig 1). Taken together, these data show that treatment with 250μM palmitate for 48hours impaired insulin secretory response but with no effect on cell viability.

### *Pex11*β knockdown decreases PMP-70 protein expression

MIN6 cells were transfected using siRNA probe s71497 against *Pex11*β. At 96hr post transfection, *Pex11*β gene expression was significantly decreased by ≥80% (Fig 1A) with compared with the scrambled negative control (P<0.001). PMP-70 is a major component of the peroxisome membrane and is used to measure peroxisome abundance (17, 29). At 96hrs post-transfection, there was a 30% decrease in PMP-70 expression relative β-actin (Fig 1B) in the *Pex11*β knockdown cells compared with the scrambled controls (p<0.001). Fig 1C is a representative blot of PMP-70 expression 96hrs post transfection with 2 probes, s71497 and s71499. Only probe s71497 led to decreased PMP-70 expression and was therefore used for subsequent *Pex11*β knockdown experiments.

**Fig 1.**
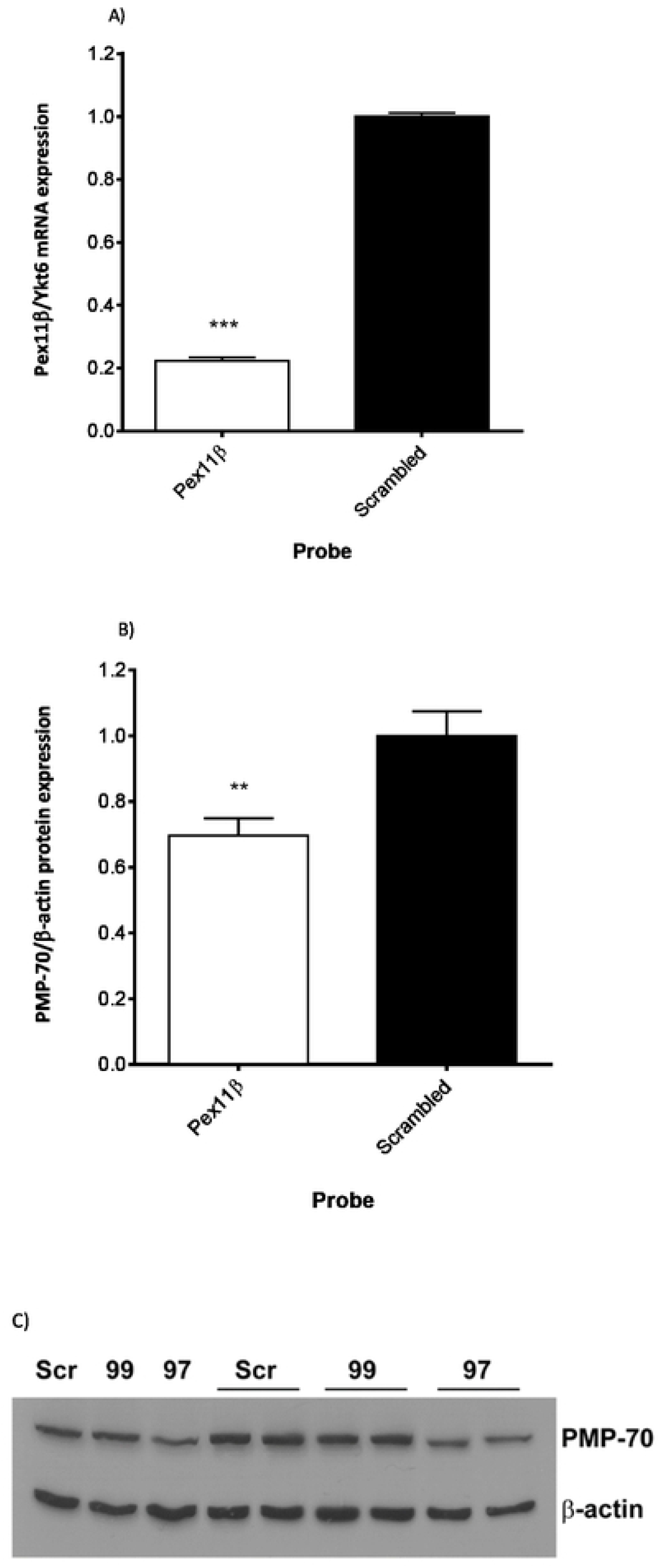
*Pex11*β knockdown and PMP-70 expression in MIN6 cells. MIN6 cells were transfected with probes (s71497) against *Pex11*β or scrambled control. **A**) *Pex11*β mRNA expression was measured after 96hrs by relative quantification using the reference gene Ykt6. The graph shows the mean ± SEM for 3 separate experiments carried out for 3 separate transfections (n=9), *** p<0.001. **B**) PMP-70 protein expression relative to β-actin expression was assessed at 96 hrs following *Pex11*β mRNA knockdown. The graph shows the mean ± SEM for 3 separate experiments were carried out for 2 separate transfections (n=6), **p<0.01. For both **A)** and **B)**, data are normalised to the scrambled control. **C)** A representative blot of PMP-70 and β-actin expression 96 hrs post-transfection (Scr=scrambled control, 99=probe s71499 and 97=probe s71497). Decreased PMP-70 expression was not achieved with s71499, so probe s71497 was used for all subsequent experiments.

To further investigate peroxisome abundance 96hrs post-transfection, cells were seeded onto coverslips and stained for PMP-70. *Pex11*β knockdown cells had fewer peroxisomes compared with the scrambled control cells (Figs 2A to 2D), corroborating the western blotting results. Analysis and quantification of the images showed a significant decrease of PMP-70 expression in *Pex11*β knockdown cells compared with the scrambled controls (P<0.05; Fig 2E).

**Fig 2.**
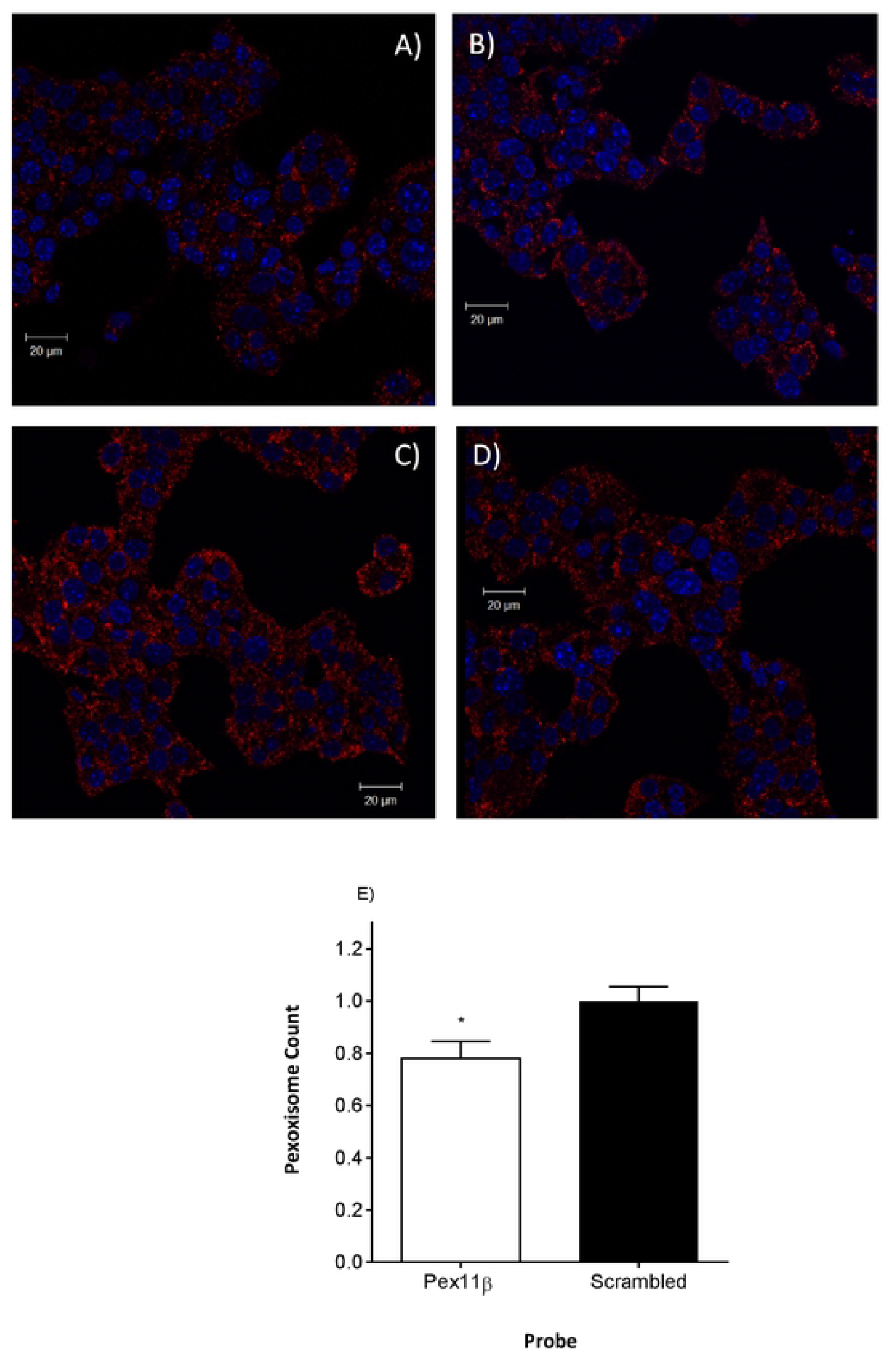
Immunofluorescent visualization of peroxisomes following *Pex11*β knockdown. MIN6 cells were stained for PMP-70 as a marker of peroxisomes (red) 96hrs after transfection with the s71497 siRNA probe against *Pex11*β. DAPI was used for nuclear counterstaining (blue). **A**) and **B**) show cells transfected with the probe against *Pex11*β, while **C**) and **D**) are cells transfected with scrambled control. For the quantification of peroxisomes **E**) 9 images from a single transfection were analysed from 4 separate transfections and the number of peroxisomes per area of the image covered by the cells was calculated. Data are normalised to scrambled control and presented as mean ± SEM, * p<0.05.

Having established that peroxisome abundance was decreased 96 hrs post *Pex11*β knockdown, all subsequent experiments followed the same design. 48hrs post transfection, MIN6 cells were incubated with either 250μM palmitate or BSA control for a further 48hrs. At 96 hrs post transfection, the relevant measure was made (eg ROS generation, GSIS).

### *Pex11*β knockdown does not alter mitochondrial respiratory function

Oxygen consumption rate (OCR) was measured in MIN6 cells following siRNA transfection and treatment with palmitate using the Seahorse XF24 Analyzer (Fig 3A). In *Pex11*β knockdown cells, palmitate treatment increased ATP synthesis by oxidative phosphorylation (Fig 3B) and basal mitochondrial OCR (Fig 3C) compared with the BSA controls (both, P<0.01). The same pattern was seen in the scrambled controls treated with palmitate, with a trend for an increase in both ATP synthesis and basal OCR. However, there were no differences between the *Pex11*β knockdown and scrambled control cells for ATP synthesis or basal mitochondrial OCR in the presence of palmitate. These data indicate that mitochondrial respiratory function in the presence of palmitate was not affected by peroxisome depletion.

**Fig 3.**
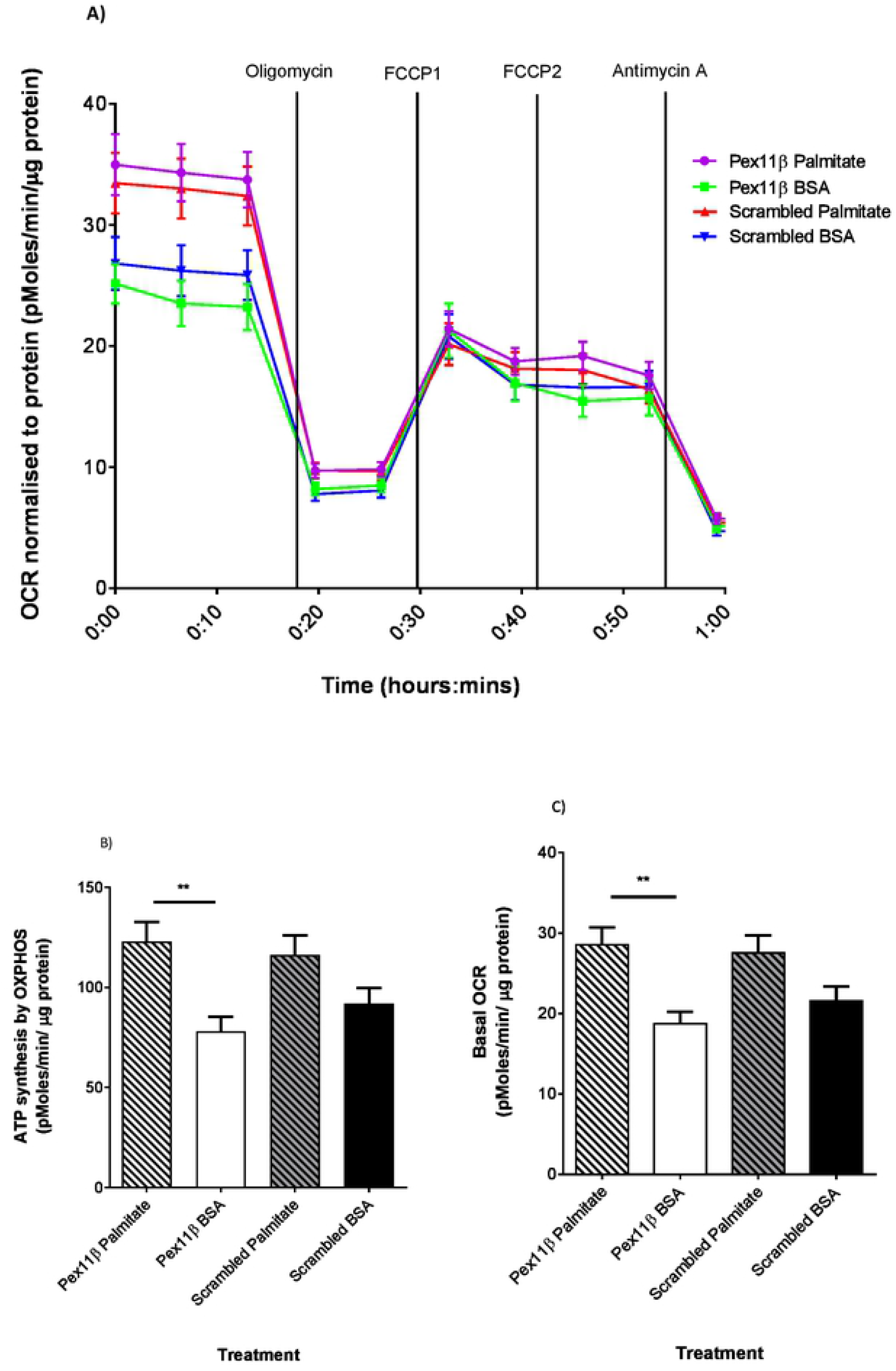
Palmitate treatment and mitochondrial respiratory function following *Pex11*β knockdown. 48hrs post transfection MIN6 cells were incubated with 250μM palmitate or BSA control for a further 48hrs. Following this, media was replaced with basic seahorse media containing palmitate and cells were incubated in a CO_2_ free incubator for 1hr prior to measuring oxygen consumption rate using the Seahorse XF24 Analyzer. In order to assess fatty acid oxidation, **A**) mitochondrial respiration was analysed by injections of compounds known to alter mitochondrial function: Oligomycin, FCCP, and Antimycin A. **B**) ATP synthesis by oxidative phosphorylation (OXPHOS) and **C**) Basal OCR were calculated as previously described. Data from 2 separate experiments each with 5 separate transfections were normalised to total protein content and expressed as pMoles/min/μg protein. Data are presented as mean±SEM. **p<0.01 palmitate vs BSA control.

### *Pex11*β knockdown decreases palmitate induced ROS production

Following treatment with 250μM palmitate or BSA control for 48hr, ROS production was measured in the transfected MIN6 cells using the dye DCFDA (Fig 4). In both *Pex11*β knockdown and scrambled control cells, palmitate significantly increased ROS production (P<0.001). In the presence of palmitate, ROS levels were markedly lower in the *Pex11*β knockdown cells compared with the scrambled controls (P<0.001).

**Fig 4.**
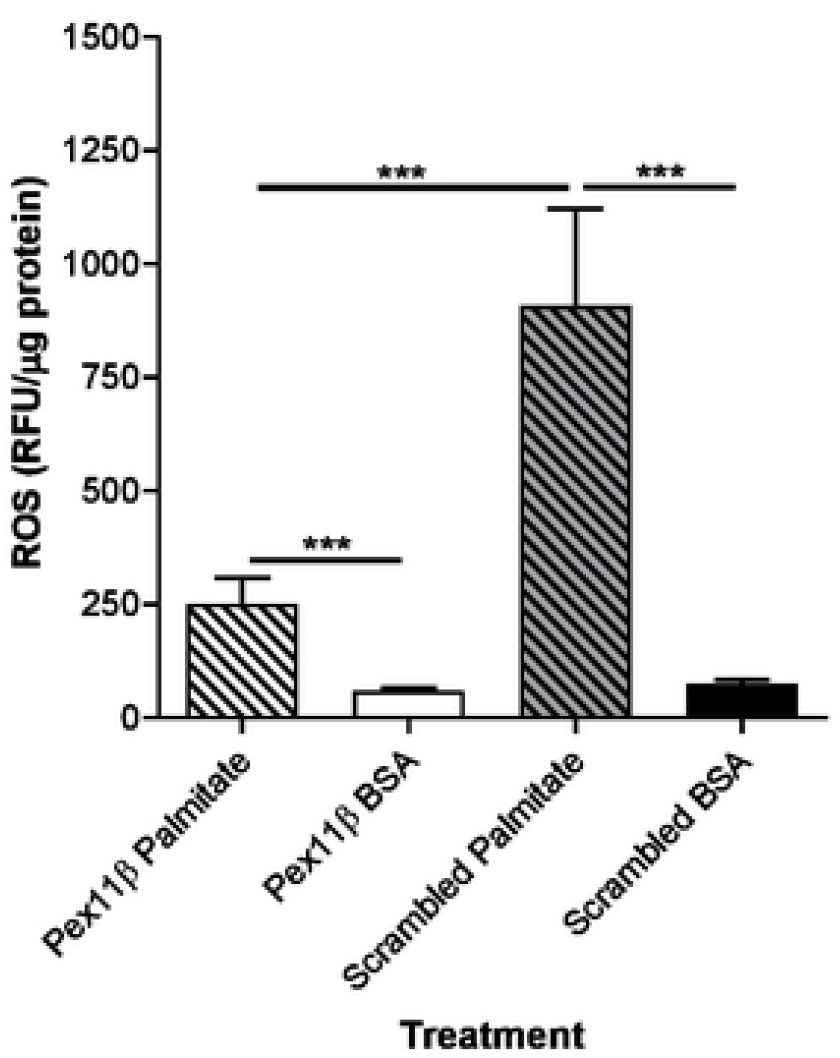
Palmitate treatment and ROS production following *Pex11*β knockdown. 48hrs post transfection MIN6 cells were treated with either 250μM palmitate or BSA for a further 48hrs in medium containing 25mM glucose. The ROS detection dye DCFDA was added to the cells and ROS determined in triplicate for each transfection. The figure shows ROS in MIN6 cells transfected with *Pex11*β knockdown or scrambled siRNA and incubated in palmitate or BSA for 48hrs. Data from 6 separate transfections are normalised to total protein content. Data are presented as mean±SEM. ***P<0.001

### *Pex11*β knockdown reverses the inhibitory effect of palmitate in GSIS

Following treatment with 250μM palmitate or a BSA control for 48hrs, GSIS was determined 96 hrs post-transfection (Fig 5A). At 25mM glucose, palmitate treatment decreased GSIS in the scrambled control cells (2.54±0.25 vs 7.07±0.83 ng/hr/μg protein; palmitate vs BSA; P<0.001). Palmitate treatment had a similar but diminished inhibitory effect in the *Pex11*β knockdown cells (4.04±0.46 vs 6.40±1.02 ng/hr/μg protein; palmitate vs BSA). In the presence of palmitate, GSIS was significantly higher in the *Pex11*β knockdown versus scrambled control cells (4.04±0.46 vs 2.54±0.25 ng insulin/hr/μg protein; P<0.05). These data show that the inhibitory effect of palmitate on GSIS was less pronounced following peroxisome depletion.

**Fig 5.**
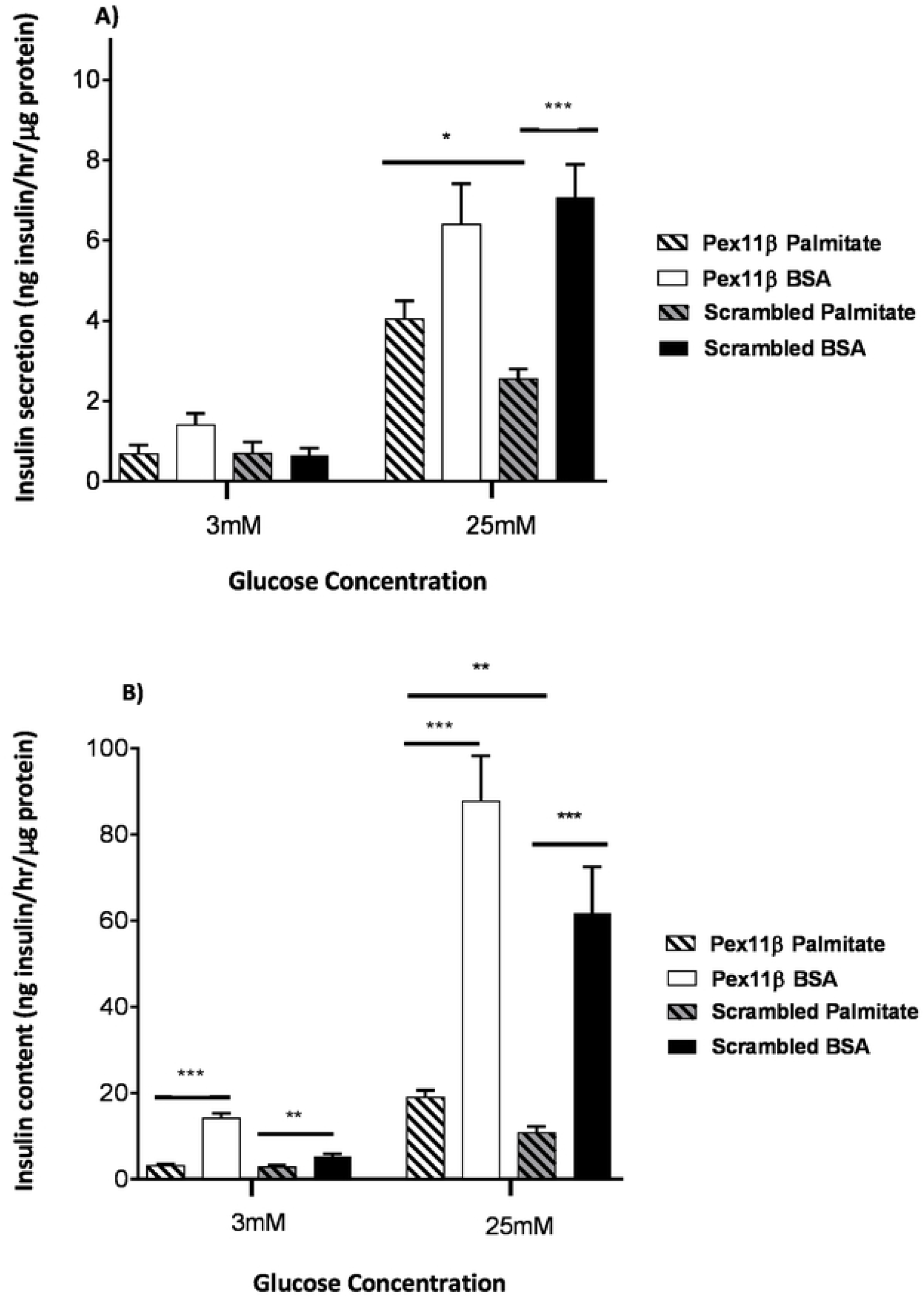
Palmitate treatment and insulin secretion and content following *Pex11*β knockdown. 48hrs post transfection, MIN6 cells were treated with 250μM palmitate or BSA control for a further 48hrs. Cells were then stimulated with either 3mM or 25mM glucose. **A**) Insulin secretion and **B**) insulin content were determined, and data from 3 separate experiments carried out in triplicate presented as mean±SEM. *p<0.05, **p<0.01, ***p<0.001

### *Pex11*β knockdown reverses the adverse effect of palmitate on insulin content

We next investigated whether *Pex11*β knockdown altered intracellular insulin content (Fig 5B). At 25mM glucose, insulin content was decreased by >80% in scrambled siRNA control cells treated with palmitate versus BSA control (10.78±1.37 and 61.67±10.78 ng insulin/μg protein, P<0.001). Similarly, palmitate treatment significantly decreased insulin content in the *Pex11*β knockdown cells (18.92±1.70 and 87.71±10.50 ng insulin/μg protein, P<0.001). However, insulin content was greater in *Pex11*β knockdown cells compared with the scrambled control cells in the presence of palmitate (18.92±1.70 vs 10.78±1.37 ng insulin/μg protein, P<0.01). These data show that the effect of palmitate to decrease insulin content was less pronounced in the *Pex11*β knockdown cells.

## Discussion

We found that *Pex11*β knockdown decreased peroxisome abundance, decreased palmitate mediated ROS generation, and reversed the inhibitory effect of palmitate on GSIS. These beneficial changes were independent of mitochondrial respiratory function which was unaffected by *Pex11*β knockdown. Our investigations have established that targeted peroxisome depletion counteracts the adverse effects of palmitate on pancreatic beta-cell secretory function.

It is well established that chronic palmitate treatment decreases GSIS (5, 7, 8, 30). Watson et al found that 48hours treatment of MIN6 cells with 400μM palmitate decreased GSIS but also increased the pro-apoptotic marker caspase 3/7 activity (8). In addition, studies using primary rat islets have shown that 500μM palmitate treatment for 48 hours was cytotoxic with around 25% of cells damaged (31). Based on these findings we elected to use 250μM palmitate treatment for 48hours, and found that under these conditions there was a clear decrease in GSIS but with no evidence of increased cell damage or apoptosis relative to the BSA control.

Both peroxisomes and mitochondria are in involved in the metabolism and oxidation of palmitate and related long-chain fatty acids (12). Peroxisome β-oxidation generates H_2_O_2_ and shortens the fatty acid chain length prior to transfer to the mitochondria for complete oxidation and ATP generation. We first explored whether *Pex11*β knock down and decreased peroxisome abundance altered mitochondrial respiratory function. As shown in Fig 3, palmitate increased basal mitochondrial OCR and ATP generation to a comparable degree in the *Pex11*β knockdown and scrambled control cells. We therefore conclude that the beneficial effects of *Pex11*β knockdown and peroxisome depletion on beta-cell function were independent of changes in mitochondrial respiratory function.

We next examined ROS generation following palmitate treatment. As shown in Fig 4, palmitate treatment increased ROS generation, but this was markedly lower in the *Pex11*β knockdown compared with the scrambled control cells. This is consistent with the observation that peroxisomes are a major source of H_2_O_2_ production in insulin producing cells (17). It has been proposed that H_2_O_2_ produced by the peroxisomes may contribute to lipotoxicity due to the low levels of catalase expressed in pancreatic beta-cells (14–16). The over-expression of catalase in peroxisomes and the cytosol was found to decrease H_2_O_2_ generated by both the peroxisomes and the mitochondria, and to protect against palmitate toxicity in a rat beta-cell line and isolated rat islets (17). However, the relative contributions of the peroxisomes and mitochondria to the ROS mediated cytotoxicity was not be defined. Through targeted *Pex11*β knockdown and peroxisome depletion we have extended these findings and identified a clear and independent role for peroxisomes in palmitate mediated pancreatic beta-cell dysfunction.

It is recognised that ROS and specifically H_2_O_2_ can adversely affect beta-cell function. Insulin secretion decreased from rat pancreatic islets when exposed to low H_2_O_2_ concentrations similar to that of physiological levels. It was found that this was through the reduction of [Ca^2+^]_i_ oscillation amplitude, which in turn inhibited GSIS (34). These observations suggest that palmitate increases H_2_O_2_ production from peroxisomes which, through reduced [Ca^2+^]_i_ oscillation amplitude, inhibit the insulin secretory response to glucose.

Previous studies have reported decreased insulin content in rodent and human pancreatic beta-cells after palmitate treatment (32, 33), with evidence that the mechanism involves decreased insulin translation (33). As shown in Fig 5B, palmitate decreased insulin content in both *Pex11*β knock down and control cells at high glucose, but this was less pronounced in the *Pex11*β knock down cells. It is not clear whether ROS and specifically H_2_O_2_ exert an inhibitory effect on insulin synthesis as well as secretion. Nonetheless, our findings show that *Pex11*β knockdown and decreased peroxisome abundance reverses the adverse effects of palmitate on both insulin secretion and insulin content.

Our results support the concept that peroxisomes are involved in lipotoxicity and pancreatic beta-cell dysfunction, and thereby might well contribute to the pathogenesis of type 2 diabetes. However, exploring peroxisomes as potential targets for therapies that counteract the impact of lipotoxicity on beta-cell function requires careful consideration. This follows a recent study that described a beta-cell specific Pex5 knockout mouse (35). Pex5 is required for the import into the peroxisome of the majority of the enzymes essential for lipid metabolism. While there was evidence of *decreased* GSIS, this appears to be a model of severe peroxisome dysfunction. First, the beta-cell specific Pex5 knockout resulted in a 5 fold increase in long chain fatty acid levels in the systemic circulation. Second, the phenotype extended beyond peroxisome dysfunction, with evidence of mitochondrial dysfunction, increased beta-cell apoptosis and decreased beta-cell mass. So while partial peroxisome depletion appears to improve beta-cell function in the presence of palmitate, there is the risk that more severe peroxisome dysfunction might trigger secondary off-target changes and an overall impairment of beta-cell function.

In conclusion, we have shown that *Pex11*β knockdown decreased peroxisome abundance, decreased palmitate mediated ROS generation, and reversed the inhibitory effect of palmitate on GSIS. This highlights a potential role of peroxisomes in pancreatic beta-cell lipotoxicity and the pathogenesis of type 2 diabetes.

## Authors Contributions

HRB, DG and MW conceived and designed the study. HRB, CT, AEB, SM, AH, AR and MvG conducted the experiments, and generated and analysed the data. All authors contributed to data interpretation and the writing of the paper.

## Supplementary Information

**S Fig 1. The effect of palmitate incubation on insulin secretion and cytotoxicity in MIN6 cells.** MIN6 cells were incubated with either 250μM palmitate or BSA control for 48hrs. **A**) Cells were challenged with either 3mM or 25mM glucose and insulin secretion was measured. Data are from 3 separate experiments carried out in triplicate are presented as mean±SEM. **B**) LDH release into the medium and compared with the LDH assay positive control. Data are from 3 separate experiments carried out in triplicate are presented as mean±SEM. **C**) Caspase 3/7 activity was measured using the luminescent Caspase-Glo^®^ 3/7 assay. Staurosporine was used as a positive control. Data are from 6 separate experiments carried out in triplicate are presented as mean±SEM. *P<0.05, ***P<0.001 vs BSA control.

